# Identification of rare pseudouridylated microRNA by comprehensive smallRNA bisulfite sequencing of mouse and human tissues

**DOI:** 10.64898/2026.05.14.725264

**Authors:** Christian Fagre, Wendy Gilbert

**Affiliations:** Department of Molecular Biophysics and Biochemistry, Yale University, New Haven, Connecticut, United States of America

## Abstract

Pseudouridine (Ψ) is an important post-transcriptional modification of many noncoding RNAs that is under-characterized in microRNA (miRNA) due to historical limitations in pseudouridine mapping methods. Ψ modification stabilizes RNA duplex structures and could therefore play an important role in miRNA target binding and repression. To investigate the extent to which mammalian miRNAs are modified with Ψ, we profiled the modification landscape of short (<30 nt) RNA in human cells and mouse tissues using bisulfite sequencing. Our approach was powered to detect small RNA pseudouridylation based on robust detection of known Ψ positions in tRNA fragments (tRFs), some of which show tissue-specific patterns of modification. In contrast with tRFs, we find that miRNA pseudouridylation is exceedingly rare, with a single modified miRNA (miR-3068-5p) identified in mouse tissues. Pseudouridylated miR-3068-5p diSerentially repressed predicted miRNA targets with less stable miRNA:mRNA pairing modes. This study fills a long-standing gap in transcriptome-wide Ψ profiling and reveals a new potential function for Ψ as a modulator of activity of small regulatory RNAs.

## Introduction

Post-transcriptional modification of RNA is a high regulated process essential to proper RNA function and processing. Pseudouridine (Ψ), an isomerization of uridine installed by pseudouridine synthases (PUS), is the most abundant RNA modification in eukaryotes. The majority of Ψ is installed in rRNA, tRNA, and snRNA, and in the last two decades a wealth of new Ψ profiling methods have uncovered modification of nearly every class of RNA. Pseudouridine can play a variety of functional roles in a given RNA, including affecting RNA stability, RNA processing, RNA secondary structure, RNA-protein interactions, and codon decoding^1–3^While maps of pseudouridylation across the eukaryotic transcriptome have been widely expanded in recent years, limitations in mapping methodologies mean these maps are still likely far from complete, and thus the scope of pseudouridine function in the cell is still not fully understood.

MicroRNAs (miRNA) are essential regulators of gene expression. These 18-22 nucleotide (nt) RNAs associate with Argonaute (AGO) proteins to form RNA-induced silencing complexes (RISC), and bind to specific mRNA transcripts by complementary pairing with 6-8 nt seed sequences, leading to translational repression or RNA degredation^4–6^ ( The production and processing of miRNA in the cell is tightly regulated and cell-type specific. A major mechanism for miRNA regulation is post-transcriptional modifications, which can include changes to the microRNA sequence through alternative processing of miRNA precursors^7^, or by 3′ and 5′ non-templated nucleotide additions^4^. MicroRNAs are also subject to installation of internal chemical modifications, such as inosine (I)^8,9^, N6-methyladenosine (m6A)^10^, and N5-methycytosine (m5C)^11^. miRNA modifications can lead to changes in miRNA stability, effectiveness at target repression, and targeting preferences^9^. Ψ is a potent stabilizer of short RNA duplexes and secondary structure^12^, so pseudouridylation of miRNA could have strong effects on target affinity and therefore miRNA function. Additionally, Ψ is demonstrated to impact RNA-protein interactions^13–15^, potentially altering functional loading into the RNA-induced silencing complex (RISC).

Pseudouridine has been discovered in nearly every class of RNA and is especially abundant in non-coding small RNA (< 200 nucleotides), such as tRNA, snRNA, and snoRNA^1^. The first transcriptome-wide maps of pseudouridine leveraged high-throughput sequencing of RNA subjected to a chemical treatment with N-cyclohexyl-N0-(2-morpholinoethyl)carbodiimide metho-p-toluenesulfonate (CMC)(Pseudo-seq^16^, CeU-seq^17^, PSI-Seq^18^, Ψ-seq^19^). The CMC treatment in these methods forms bulky chemical adducts specific to Ψ, at which reverse-transcriptase terminates during library construction, leading to truncation of reads at pseudouridine positions in the final sequencing dataset. While effective at mapping Ψ in most types of RNA, these methods likely would fail to detect miRNA pseudouridylation, as very short, truncated miRNA reads would either be depleted from the sequencing library or fail to align. Methods using anti-Ψ antibody enrichment have shown evidence of pseudouridylated miRNA populations in mouse, but this approach lacks single-nucleotide positional information or estimates of modification stoichiometry.^20^ Thus, the current Ψ landscape lacks a high-resolution map of microRNA modification.

Recently, multiple methods have been developed that allow for sensitive detection of Ψ by formation of Ψ-specific mutation signatures rather than truncations. These methods allow for full-length capture of modified RNA fragments and would in theory tolerate mapping of modified miRNAs. Most of these new methods employ sodium bisulfite and/or sodium sulfite treatments (RBS-MaP^21^, BID-Seq^22^, PRAISE^23^), which reacts with pseudouridine to form chemical adducts that are skipped by certain reverse-transcriptases, leading to a deletion signature in the final sequencing dataset. These methods have effectively mapped pseudouridine across mRNA, rRNA, snoRNA, and snRNA with single-nucleotide precision, yet none have specifically interrogated miRNA populations.

Here we sought to chart the Ψ modification landscape of mammalian microRNAs and other small RNAs such as tRNA fragments and piRNAs with single-nucleotide resolution using BID-Seq. We establish that this method is able to accurately detect tRNA Ψ sites in tRNA fragments from human cells and identify tissue specific differences in tRNA fragment modification across mouse tissues. Across microRNA populations in mouse and human, we identify a single modified miRNA, miR-3068-5p, as present and highly modified across four mouse tissues. By mRNA-sequencing of cells transfected with pseudouridylated miR-3068-5p, we find evidence that modification can enhance repression of specific targets, suggesting regulatory potential for small RNA modification.

## Results

### Bisulfite-based pseudouridine profiling successfully identifies pseudouridine in small RNA populations

We first compared chemical probing and reverse-transcription conditions to identify one most suitable to detecting smallRNA modification. Specifically, we sought conditions leading to high pseudouridine-specific mutation signatures and high processivity past adducted Ψ sites. With a pool of reporter RNAs containing a single 100% U or Ψ position flanked by random nucleotides [S. Fig. 1A], we found that CMC treatment could generate a Ψ-specific mutation signature upon reverse transcription with SuperScript II and 6 mM MnCl_2_ [Fig. S1B]. However, this signature was highly variable across sequence contexts and would therefore be unable to comprehensively identify Ψ positions. Using sodium bisulfite with the same reverse-transcription conditions (SuperScript II, 6 mM MnCl_2_) we observed a 1-2 nt Ψ-specific deletion signature that was more penetrant (up to 80% deletion rates) and consistent across sequence contexts [Fig. S1E]. We additionally tested Marathon RT, a highly processive group II intron reverse transcriptase, resulted in a consistent T-C mutation signature at Ψ, however with a weaker penetrance than SuperScript II + MnCl_2_ [Fig S1G]. Finally, we tested conditions used in the BID-Seq method^22^, included adjusting the pH of bisulfite treatment to 7 from 5, and using SuperScript IV for reverse transcription.

**Figure 1:**
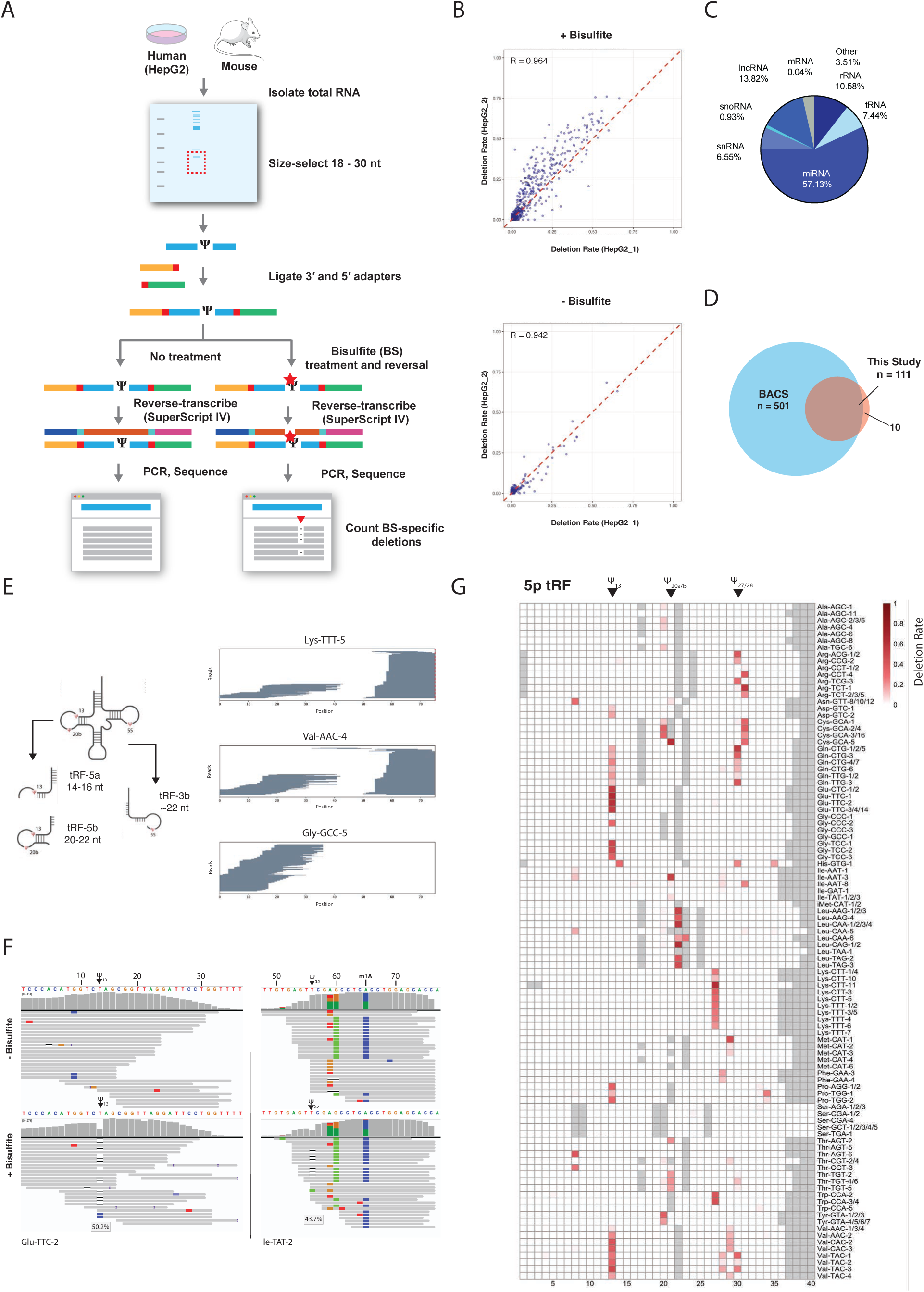
tRNA fragment pseudouridine profiling in HepG2 cells A: Schematic of smallRNA BID-Seq. Small RNA was extracted from HepG2 human cells lines or mouse tissues (lung, cerebellum, lung, testes) and subjected to BID-Seq^22^. **B.** Correlation rates of deletion rates per-position across mapped tRNAs in untreated and bisulfite treated samples. R = Pearson’s R. **C**: Alignment rates of smRNA libraries to different RNA classes. **D**. Venn diagram of overlap in tRNA Ψ site-calling between this method and BACS^28^. Correlation rates of deletion rates across mapped tRNA fragments in untreated and bisulfite treated samples. R = Pearson’s R. **E**: (Left) Cartoon of tRNA fragment (tRF) biogenesis. (Right) Example pileups of reads mapping to full-length tRNAs, showing distinct fragment boundaries. **F.** Example IGV browser views of untreated (top) or bisulfite-treated samples at two tRNA sites. Called Ψ position indicated with arrow. **G**: Heatmap of deletion rates for the top 100 expressed 3′-tRFs, grouped by isodecoders.

These conditions led to a 1 nt deletion signature that was more consistent across sequence contexts [Fig. S1H]. Additionally, the higher pH treatment avoided global C->T conversions normally caused by bisulfite treatment, which we predicted would allow for more accurate mapping of smallRNAs. For these reasons, we proceeded with the conditions of BID-Seq for our smallRNA Ψ profiling study.

To map pseudouridylation in small RNA fractions, we size-selected the fraction of 18-30 nt small RNAs (smRNA) from a human hepatocellular carcinoma cell line (HepG2) and performed Ψ profiling [Fig. 1A]. After alignment to the human transcriptome, we observed clean 1-nt deletion signatures specific to bisulfite-treated samples with excellent reproducibility between biological replicates (R = 0.942) [Fig 1B]. The majority of reads captured in these libraries mapped to microRNA (57%), with smaller fractions mapping to tRNA, (7.4%) snRNA (6.5%) and snoRNA (0.9%) [Fig. 1C]. Since tRNA are highly modified with Ψ at known positions at high stoichiometry, we first examined tRNA-mapping reads to confirm the sensitivity of our approach to detect Ψ in ∼22-nt RNA species. We first compared positions of bisulfite-specific deletions in tRNA-aligned reads to pseudouridine sites identified through profiling of full-length tRNA by the chemically orthogonal 2-bromoacrylamide based mapping method BACS^28^. Overall, of the 111 Ψ sites we identified in this study, 103 are reproduced in BACS [Fig. 1D]. Of the non-overlapping sites, 4 are immediately adjacent to a site reported in BACS within a UU or UUU context where deletion positions cannot be accurately resolved, and 3 are in tRNAs not present in the BACS dataset but at common Ψ positions (13, 27, 55). In these libraries, we observed an enrichment of read alignments to tRNA 3′ or 5′ ends, consistent with processed tRF-3 or tRF-5 species rather than degraded tRNA, consistent with processed tRF-3 or tRF-5 species rather than degraded tRNA [Fig. 1E].^24^ In order to reduce multimapping of these fragments to distinct tRNAs with homologous 3ʹ or 5ʹ ends, we built a tRF fragment reference consisting of the first or last 35 nt of all tRNAs, with identical fragments collapsed into a single reference. After re-aligning to this reference set, we reproducibly captured 173 3′ fragments and 187 5ʹ unique fragments representing all 49 tRNA families [Fig. S2C-D]. These deletion signatures in tRNA fragments occurred at known commonly modified uridines, such as U_13_ in Glu-TTC-2, and U_55_ in Ile-TAT-2, with less than 0.1% deletion rates in untreated samples [Fig. 1G]. From these data we conclude that we can accurately identify Ψ in RNA in similar length to miRNA.

Looking across all tRFs, we captured extensive modification across 5′ tRFs at position U_20a/b_, U_27_, and U_28_. While we were able to detect U_55_ modification in many tRFs [Fig. S2E], the vast majority of captured 3′ fragments start at or after position 55, leaving too sparse coverage for accurate detection (for example, Lys-TTT-5 and Val-AAC-4 [Fig. 1C]). In addition to Ψ, we detected mutation signatures from other tRNA modifications: m^1^G (G_7_) appears as a bisulfite-independent deletion [Fig. S2B], and m^1^A (A_58_) appears as an A – T mutation, a signature observed with other highly processive reverse-transcriptases when profiling tRNA^29,30^ [Fig. 1D]. BID-Seq therefore may be compatible with other tRNA-profiling methods such as mim-tRNA-seq^30^ for more comprehensive characterization of tRNA and tRF modifications.

### tRNA fragment profiling in mouse show tissue-specific patterns of pseudouridylation

We similarly profiled tRF Ψ in small RNA isolated from mouse lung, liver, testes, and cerebellum of three mice per tissue. Similar to human samples, in lung, liver, and cerebellum roughly 10-20% of reads mapped to tRNA allowing for sufficient coverage for Ψ profiling [Fig. 2A]. In testes, the vast majority of reads (84.5%) mapped to piRNA loci. This is expected due to the high levels of piRNA in testes, but these samples were excluded from downstream tRF profiling due to poor tRNA coverage (2.9%). We captured distinct alignment patterns consistent with processed tRFs and observe a wider spread of tRF-3 lengths, allowing for sufficient coverage for confident quantification of Ψ_55_ across a larger set of tRFs [Fig. S3B,D]. We observed consistent bisulfite-dependent deletion peaks at position 55 across all tissues, and tRNA-specific peaks at positions 13, 20/20a/20b, 50, and 67 [Fig. 2B].

**Figure 2:**
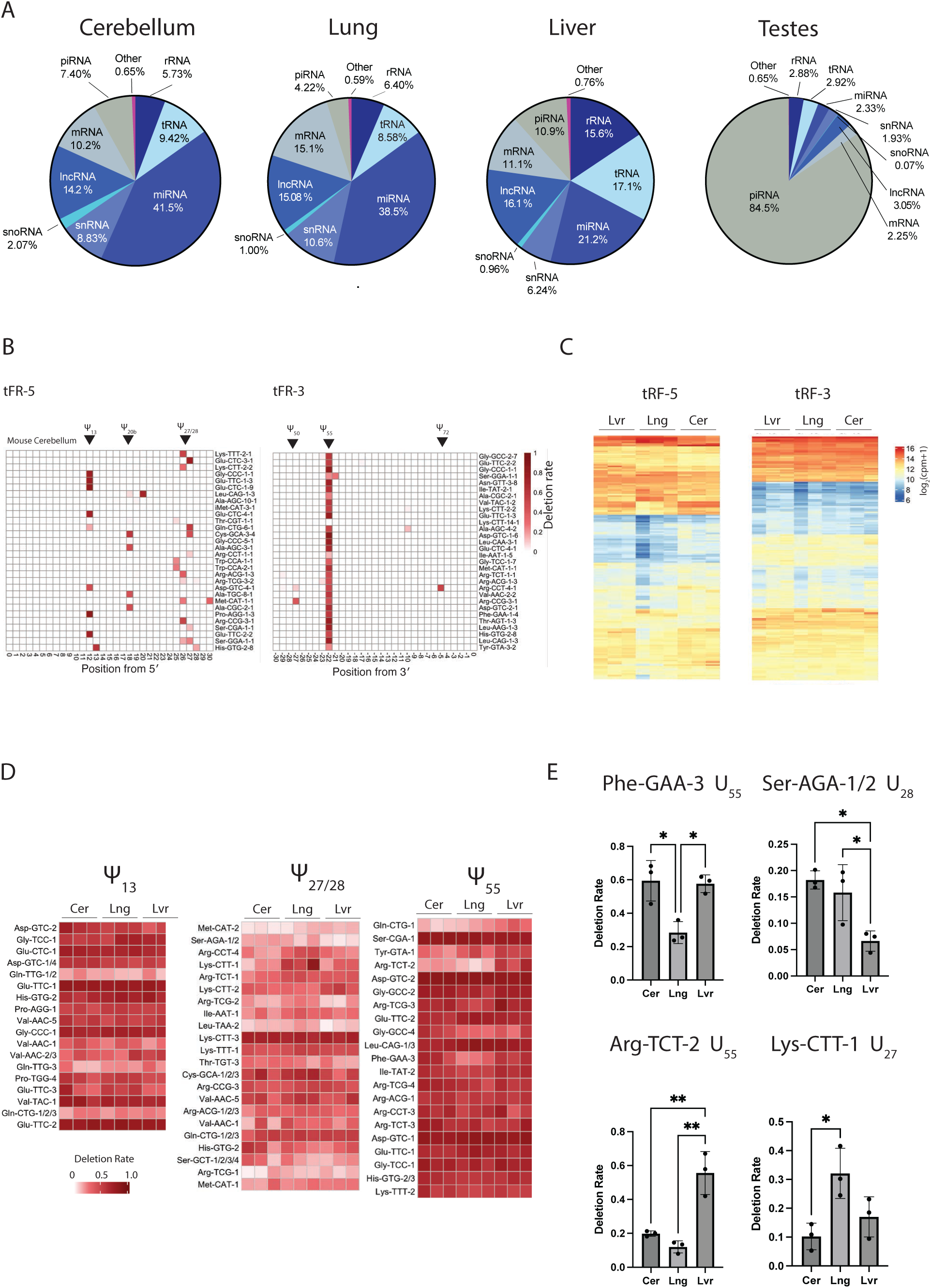
Cross-tissue mouse profiling of tRNA fragment modifications. **A**. Alignment rates of smRNA libraries to different RNAs. **B**. Example pileups of reads mapping to full-length tRNAs, showing distinct fragment boundaries. **C**. Heatmap of deletion rates for 3p and 5p tRFs in mouse cerebellum. **D**. Clustered Heatmaps of normalized tRF abundance across mouse liver, lung, and cerebellum. **E**. Heatmaps of deletion rates at specific called Ψ sites across tissue replicates. **F**. Barplots of deletion rates of sites in specific tRFs. One-way ANOVA, * = p < 0.05 ** = p < 0.01.

There is evidence that certain tRNA modifications, such as acp^3^U, have tissue-specific modification patterns^31^, and we were curious if this was the case with tRFs captured in our dataset. First, we determined whether patterns of tRF abundance differed across tissue contexts, which could result from tissue-specific tRNA expression or processing. Largely, tRF patterns were similar across lung, cerebellum, and liver [Fig 2C], with high pairwise correlations between all samples (R > 0.78) (Fig. S3A). Likewise, patterns of pseudouridylation were mostly identical [Fig. 2D], with a few notable tissue-specific differences [Fig 2E]. In Phe-GAA-3, we detected Ψ_55_ as ∼50% less modified in lung compared to cerebellum and liver. In addition, Ψ_55_ in Arg-TCT-2 showed significantly higher modification in liver compared to cerebellum and lung (3-fold and 5-fold, respectively). In 5′ fragments, we observed significant modification differences at U_27_ / U_28_ in Ser-AGA-1/2 and Lys-CTT-1. These results suggest the potential for tissue-specific pseudouridylation to affect regulatory functions of tRFs.^32,33^ It was previously reported that the pseudouridylation status of U_8_ of 5′ tRFs derived from Ala/Cys/Val tRNAs impact their function as regulatory RNAs^34^. We did not observe modification of these fragments in our dataset. Taken together, these profiles reveal a broad landscape of mammalian tRF pseudouridylation. The robust detection of Ψ in tRFs demonstrates the potential of this approach to uncover Ψ modifications in other RNAs of this size, such as miRNA.

### Pseudouridine is present but rare in mammalian miRNA

We next searched these small RNA libraries for evidence of Ψ modification at annotated miRNA. Bisulfite and untreated HepG2 smRNA libraries were aligned to all annotated human miRNA hairpins, and reads were analyzed for bisulfite-dependent deletion signatures. We captured 222 hairpins that met the read-depth cutoff for calling Ψ sites (100 reads) in both treated and untreated samples. [Fig. 3A]. Of these microRNAs we failed to detect any significant deletion signatures, indicating this miRNA population has little to no Ψ.

**Figure 3.**
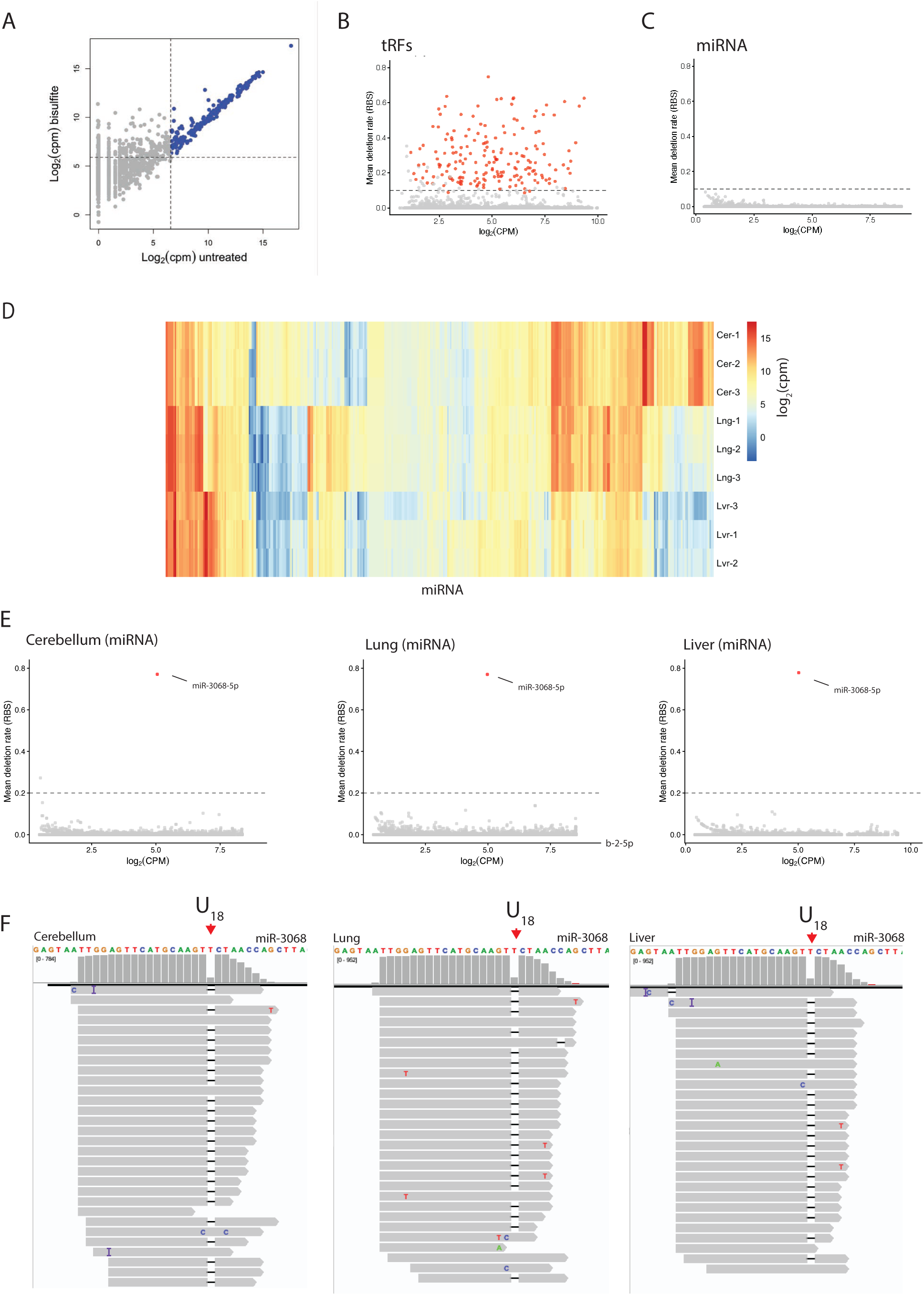
Identification of Ψ modified microRNA in human HepG2 cells and mouse tissues. **A**. Scatterplot of miRNA counts from untreated vs bisulfite-treated RNA. **B**. Scatterplots of mean deletion rates for every U in sequenced miRNA vs depth. Filtered for U’s with deletion rates > 0.01 in untreated control. **C**. Clustered heatmap of miRNA counts across all tissue replicates. **D**. Scatterplots of mean deletion rates for every U in sequenced miRNA vs depth in mouse tissues. Filtered for U’s with deletion rates > 0.01 in untreated control. **E**. Example IGV browser views of miR-3068-5p in bisulfite-treated samples. Called Ψ position indicated with arrow.

For a more comprehensive search of miRNA populations, we analyzed mouse miRNA from cerebellum, lung, and liver. As miRNAs are often expressed in a highly tissue-specific manner, we expected profiling diverse tissues would cover a greater total number of miRNAs. Indeed, we captured distinct miRNA and observed tissue-specific miRNA expression across these three tissues [Fig. 3D]. From these samples, we obtained sufficient coverage ( > 100 reads) to call modifications in 60% of annotated mouse miRNAs (746 / 1234) [Fig S4B].

In addition to providing a greater diversity of miRNA species, mouse cerebellum and liver have been demonstrated to have several fold more total pseudouridine sites in mRNA relative to common human cell lines^22^. Although the basis for this difference remains to be determined, the finding is consistent with greater activity of pseudouridine synthases towards non-canonical targets in normal tissues compared to transformed cells. Strikingly, in each of the profiled tissues we detected a single bisulfite-dependent deletion peak in miR-3068-5p at position 18, and little to no deletion signatures in any other miRNA [Fig 3D]. This deletion is present in 75-77% of reads mapping to miR-3068-5p, consistent with high to complete Ψ occupancy [Fig. 3E]. The modified uridine occurs within the sequence AGU<u>U</u>CUAACC, which matches the consensus for the TRUB1 pseudouridine synthase^35,36^ in tRNA and mRNA targets [Fig. S4D]. Overall, we conclude that in these mouse tissues miRNA pseudouridylation is exceedingly rare, as is the case in human HepG2 cells, with the single exception of highly modified miR-3068-5p.

### Pseudouridylation of miR-3068-5p alters target specificity transcriptome-wide

Pseudouridine has been shown to stabilize short RNA duplexes^12^ suggesting the potential to affect miRNA-mRNA interactions. We therefore examined whether pseudouridylation of miR-3068-5p affects mRNA targeting and repression. We synthesized miRNA duplexes with either the unmodified miR3068-5p sequence (mir3068-5p-U), or with a Ψ at position 18 (mir3068-5p-Ψ). We also synthesized control miRNAs with scrambled miR-3068-5p sequence, with or without a single Ψ (Scramble-U and Scramble-Ψ, respectively). These miRNAs were transfected into NIH/3T3 mouse fibroblast cells, and after 24 hours RNA was extracted and prepared for mRNA sequencing [Fig. 4A]. As miR-3068-5p is a poorly characterized miRNA, with no reported global analysis of its target preferences, we first examined the transcriptome-wide effects of unmodified miR-3068-5p. Our results confirm targeting of transcripts with predicted seed site matches to miR-3068-5p (TargetScan)^37^.

**Figure 4.**
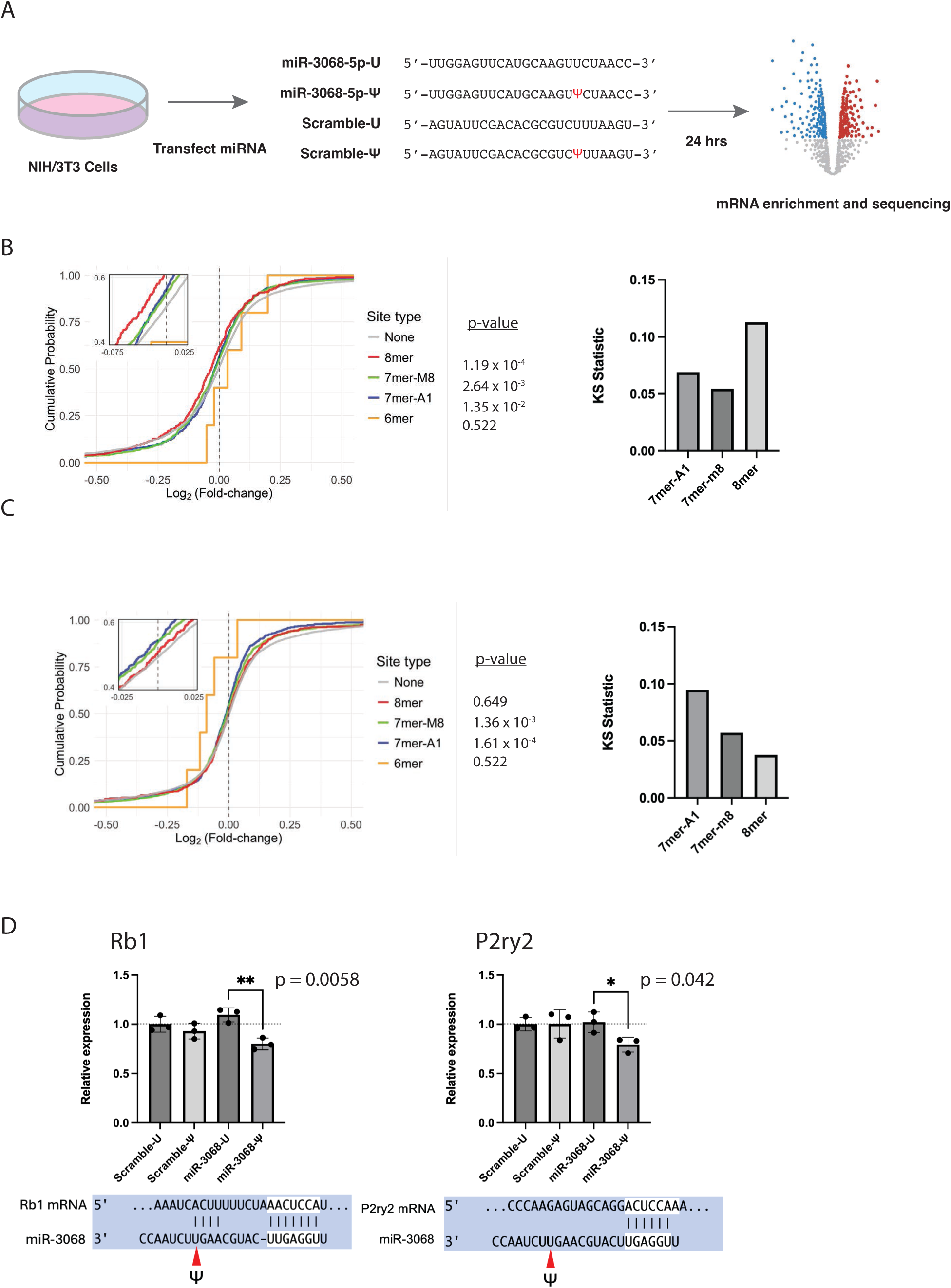
miR-3068-5p modification impacts transcript repression. **A**. Cartoon of experimental workflow. **B**. Cumulative distribution of Log_2_FC of miR3068-5p vs Scramble-U. Transcripts are binned by the strongest predicted miR-3068 site present (8 mer > 7mer-M8 > 7mer-A1 > 6mer > none) according to TargetScan. P-values calculated by Kolmogorov–Smirnov test between respective site bin and transcripts with no predicted site. **C**. Same as [B] but comparing miR-3068-5p-U to miR-3068-5p-Ψ. **D.** Relative expression of Rb1 or P2ry2 after transfection with scrmb-U, scrmb-Ψ, mir-3068-5p-U, mir-3068-5p-Ψ. B. Schematic of predicted miR-3068-5p pairing to Rb1 vs P2ry2, with Ψ position highlighted.

Grouping transcripts by class of predicted miRNA-target interaction showed that mRNA with 8mer, 7mer-m8, or 7mer-A1 sites were significantly more likely to be downregulated upon miR-3068-5p transfection than mRNA with no predicted sites. The magnitude of fold-change relative to non-target transcripts correlated with the predicted strength of each type of site^37^, with 8mer sites having the strongest shift, followed by 7mer-m8, then 7mer-A1 [Fig. 4B].

Next, we examined the effect of Ψ modification on miR-3068-5p targeting. Like the unmodified miRNA, miR-3068-5p-Ψ transfected cells showed significant repression of predicted miR-3068-5p targets, where the distribution of transcript expression fold-change shifts according to the strength of the target [Fig. S5A]. When we directly compared the fold-change repression between miR-3068-5p-Ψ and miR-3068-5p-U on a transcriptome-wide scale, we observed that miR-3068-5p-Ψ has a significantly stronger repressive effect on targets with 7mer-A1 (p = 1.61 x 10^-4^) and 7mer-M8 sites (p = 1.36 x 10^-3^) [Fig. 4C]. 6mer sites showed the greatest shift, albeit with low significance due to the small number of predicted 6mer sites included in TargetScan (p = 0.15). Notably, there was no significant difference in repression between targets with 8mer sites (p = 0.645). These results suggest that pseudouridylation of miR-3068-5p improves pairing and repression of lower strength miRNA sites (7mer and 6mer) but has no effect on targeting of stronger 8mer sites.

From these RNA sequencing experiments, we identified several transcripts that are uniquely repressed by miR-3068-5p-Ψ. P2ry2 is repressed ∼20% by miR-3068-5p-Ψ and is not significantly affected by miR-3068-5p-U [Fig. 4D]. Rb1 similarly is repressed ∼20% specifically by miR-3068-5p-Ψ. Both transcripts have predicted 7mer miR-3068-5p targets, which often engage in supplemental pairing with miRNA to improve targeting efficiency.^38^ Indeed, for Rb1 there is predicted 3′ pairing between miRNA bases 14 – 17 [Fig. 4E]. ^37^ The presence of Ψ at position 18 may strengthen this supplemental pairing, allowing for increased targeting efficiency^39^. Together, these results show that Ψ affects target repression by a mammalian miRNA and suggest the potential for Ψ to affect targets of other small RNAs that form silencing complexes with Argonaut proteins^32,33^.

## Discussion

High-throughput methods to map pseudouridine modifications have revealed abundant Ψ in diverse classes of RNA with the notable exception of small regulatory RNAs. In this study, we establish a profiling approach based on BID-Seq^22^ as an effective high-resolution method for probing very short RNA species for Ψ. Using this method, we identify a rich landscape of pseudouridylated tRNA fragments (tRFs) in human cells and mouse tissues.

In contrast to abundant Ψ in tRFs, we detected a single highly pseudouridylated miRNA, miR-3068-5p, among 968 miRNAs profiled across mouse and human samples. The presence of Ψ at position 18 of miR-3068-5p affected target repression based on RNA-sequencing of miRNA-transfected mouse fibroblasts. Given evidence that some tRFs associate with Ago to direct repression of target RNAs^32,33^, our results highlight the potential for tissue-specific tRF Ψ to regulate gene expression.

Our results demonstrate a viable strategy for profiling Ψ in tRFs, miRNA, and other small regulatory RNAs. Nevertheless, comprehensive small RNA profiling remains challenging, and BID-Seq or other bisulfite-based approaches are not perfectly suited for this purpose. For one reason, deletion signatures are not precise or accurate in all sequence contexts, especially poly-U regions ^23^. Additionally, deletions at the 3′ or 5′ terminus would manifest as a shortened read, rather than a deletion, which would not be detected as a Ψ modification in the current pipeline. The more recently developed method BACS (2-bromoacrylamide-assisted cyclization sequencing) creates U->C mutation signatures specific to pseudouridine, which can be accurately identified in any sequence context^28^. Anti-Ψ immunoprecipitation has also been used to identify modified miRNAs by enrichment^20^, however this approach lacks nucleotide resolution and pseudouridine antibodies have not been systematically evaluated for specificity across different sequence contexts. Pairing antibody-enrichment methods with mutation-mapping would alleviate all these concerns, while allowing for sensitive detection of low-abundance or lowly-modified species.

In human and mouse miRNA, across diverse tissues, we observed a single confidently modified miRNA species, miR-3068-5p, which we estimate is close to 100% modified. The lack of pseudouridylation across the broader miRNA population is surprising given the high modification rates of other small RNA species such as snoRNA, snRNA, and tRNA. Even compared to the sparse rate of mRNA pseudouridylation, this can be considered low. In mouse cerebellum, for example, 0.5% of uridine positions in examined mRNA are modified^22^ [6617 / 1,264,925 positions with coverage]. This is equivalent to 20 uridines in the much smaller miRNA sequence space, but only one was identified in this study.

One hypothesis for the paucity of Ψ in miRNA is that pseudouridylation enzymes are occluded from pri-miRNA hairpins by the microRNA processing machinery, DROSHA-DGCR8. The pseudouridine synthases responsible for many mRNA modifications (PUS1, Trub1) require a hairpin structure^35,40^, and similarly the DROSHA-DGCR8 complex recognizes a hairpin for cleaving pri-miRNA into pre-miRNA^41^. After this initial co-transcriptional cleavage^42^, miRNA sequences are bound by proteins such as Dicer and, subsequently, Argonaut, throughout the miRNA lifetime. Thus, PUS enzymes likely have a very brief window to bind and modify pri-miRNA hairpin structures.

In contrast to miRNAs processed by DROSHA-DGCR8, miR-3068-5p is one of the rare functional microRNAs that overlap an annotated snoRNA, and is presumed to be processed out of a parent snoRNA precursor, as is the case with other snoRNA-derived RNAs^43^. The majority of human snoRNA are spliced out of the introns of longer host pre-mRNAs. Given evidence that multiple pseudouridine synthases modify introns co-transcriptionally^44^,snoRNA processing may be more compatible with pseudouridylation. Further pseudouridine profiling of snoRNA populations and miRNA or snoRNA precursors is needed to illuminate the specific biogenesis pathways and enzymatic timing of this lone miRNA modification and may help explain the absence of modification in other miRNAs.

We observed that pseudouridylated miR-3068-5p had subtly different effects on mRNA targeting compared to unmodified, with weaker seed pairing sites (6-mer, 7-mer) tending to be more sensitive to Ψ modification. miR-3068-5p is modified outside the canonical seed binding region (2-8), and so there is likely no direct impact on seed site target recognition. The Ψ in miR-3068-5p is at position 18, which is often within the supplementary 3′ pairing window that can stabilize miRNA:mRNA interactions with weaker 5′ pairing^45,46^. Since one of the most strongly impacted mRNA targets, Rb1, has predicted 3′ miRNA pairing overlapping the Ψ position, we hypothesize that Ψ has a relatively strong impact on this short pairing interaction, leading to greater overall stabilization of target recognition. While miR-3068-5p is equally modified in the three tissues sequenced, our findings raise the possibility of regulated Ψ modification serving to tune the ability of miRNA-like species to repress specific targets. Such target-switching has been observed for other miRNA modifications; for example installation of a single inosine by ADAR2 into the miRNA seed site can shift miRNA-mediated repression from one set of targets to another^9^.

Many tRFs have the ability to repress gene expression in an miRNA-like manner, and are commonly associated with Argonaut proteins, suggesting they repress targets by a similar mechanism^32,33^. In HepG2 cells we captured 3′ tRFs with variable levels of Ψ_55_, which is positioned in a potential seed site for a 22nt tRF-3b fragment acting as a miRNA. Across the four mouse tissues profiled here, we observed tissue-specific Ψ modification levels for several tRFs. Our data do not include full-length tRNA sequencing and cannot differentiate tissue-specific patterns of tRNA modification^29^ from tissue-specific tRF processing.

However, it has been widely observed that tRNA modifications can protect mature tRNAs from cleavage^26,27,47,48^, thereby altering tissue-specific abundance of tRFs. It warrants further investigation whether the observed changes in tRF Ψ levels are related to modification-dependent changes to tRNA processing. Whatever the biogenesis mechanism, tissue-specific differences in tRF regulatory activity are a likely consequence of variable Ψ modification levels. The approach described here provides a framework for analyzing pseudouridylation of small regulatory RNAs to reveal undiscovered functional consequences of Ψ on gene regulation.

## Methods

### Cell culture

Human hepatocellular carcinoma cells HepG2 and mouse fibroblast NIH/3T3 cells were cultured in DMEM supplemented with 10% fetal bovine serum. Cells were grown at 37C with 5% CO2 and maintained at subconfluency.

### miRNA transfection

Sense and passenger-strand oligos (Horizon Biosciences) were annealed at 10 uM in RNA annealing buffer (30 mM HEPES, pH 7.5, and 100 mM potassium acetate) by heating to 95C for 2 minutes, then leaving to cool to room-temperature on the benchtop. NIH/3T3 cells were plated in 6-well plates at 300K cells/well 24 hours before transfection. At 80% confluence, 136 pmol miRNA duplex was transfected into each well (Trans-IT siQuest, Mirus Bio) in triplicate. 24 hours later, cells were harvested, and total RNA was extracted and submitted for mRNA sequencing at the Yale Center for Genomic Analysis.

### smRNA BID-Seq

Total RNA was Trizol-extracted from HepG2 cells, or mouse tissues (Male, C57BL/6J, 7-14 weeks, The Jackson Laboratory). Total RNA was size-fractionated using Zymo RNA Clean and Concentrator Kit, using the smallRNA protocol to enrich for < 200 nt species. smRNA fraction was further purified on a 15% Urea-PAGE gel, size-selecting 18-30 nt, eluting overnight in RNA elution buffer, and ethanol precipitating. RNA was then ligated to a pre-adenylated 3′ adapter (4.5 uL RNA, 1 uL 10 uM oCF198, 1 uL 10x T4 buffer (NEB), 2 uL PEG 8000 (NEB), 0.5 uL RNasein Plus (Promega), 1 uL T4 RNA ligase 2 truncated, K227Q (NEB)) at 22C for 16 hours. Ligated RNA was PAGE-purified away from unligated RNA/adapter. RNA was ligated to 5′ RNA adapter (oCF186) (3.5 uL RNA, 10.76 uL 10 uM oCF186, 1 uL 10x T4 buffer (NEB), 0.24 uL 10 mM ATP, 0.5 uL RNasein Plus (Promega), 1 uL T4 RNA Ligase I (NEB)) at 22C for 16 hours. Ligations were purified with MyOne Silane beads (Invitrogen).

Ligated RNA was then subject to bisulfite treatment and reversal, per BID-Seq protocol^49^, with minor modifications, and with 1/3 reserved as no treatment control. Briefly, RNA was incubated in 2.4 M Na2SO3 and 0.36 M NaHSO3 at 70C for 3 hours. RNA was desalted on MicroSpin G-25 columns and eluted in 100 uL Milli-Q H_2_O. Samples were then mixed with 100 uL 2M Tris-HCl, pH 9.0, incubated at 37C for 2 hours, ethanol-precipitated, and resuspended in 5 uL Milli-Q H_2_O. For reverse-transcription, all samples including no-treatment controls were snap annealed to RT primer (oCF180) by combining 5 uL sample, 1 uL 10 uM primer, heating to 65C for 5 minutes, and snap-cooling on ice 2 minutes. Samples were reverse-transcribed with SuperScript IV (Invitrogen) (6 uL RNA/Primer, 2 uL 10 mM dNTPs, 1 uL 100 mM DTT, 0.5 uL RNasein, 5.5 uL Milli-Q H_2_O, 1 uL SuperScript IV) at 50C for 1 hour. RT reactions were cleaned on MyOne Silane beads (Invitrogen), PCR-amplified with Truseq indexing primers, PAGE-purified, and submitted for 2×150 paired-end Illumina sequencing (Novaseq) at the Yale Center for Genome Analysis.

### smRNA read processing and alignment

FASTQ files (R1) were trimmed for adapter sequences and size-filtered with cutadapt (-a GATCGGAAGAGCACACGTCTGAACTCCAGTCA -m 17 -M 100). PCR duplicates were removed with BBMap clumpify (reorder=t, dedupe=t). UMIs were trimmed with BBMap reformat.sh (forcetrimright2=9). Reads were then sequentially aligned to RNA class references from NCBI (mouse: GRCm39, human: GRCh38) using segemehl^50^ with the -D 1 flag: ribosomal RNA > tRNA > ncRNA. Additional unaligned reads were aligned to the full m38 or h38 genome with Bowtie (--local). Deletion rates for aligned reads were calculated using SAMtools pileup and custom python scripts.

### mRNA read processing and alignment

Raw reads were trimmed with BBMap bbduk.sh using reference truseq adapter sequences (ktrim=r k=33 hdist=1). Trimmed reads were aligned to the mouse genome (m39) using STAR (--outFilterMultimapNmax 20 --alignSJoverhangMin 8 --alignSJDBoverhangMin 1 -- outFilterMismatchNmax 999 --alignIntronMin 20 --alignIntronMax 1000000 --alignMatesGapMax 1000000 --alignEndsType EndToEnd). Reads were counted using HTSeq htseq-count, and expression changes analyzed using DESeq and custom R scripts.

**Figure S1:**
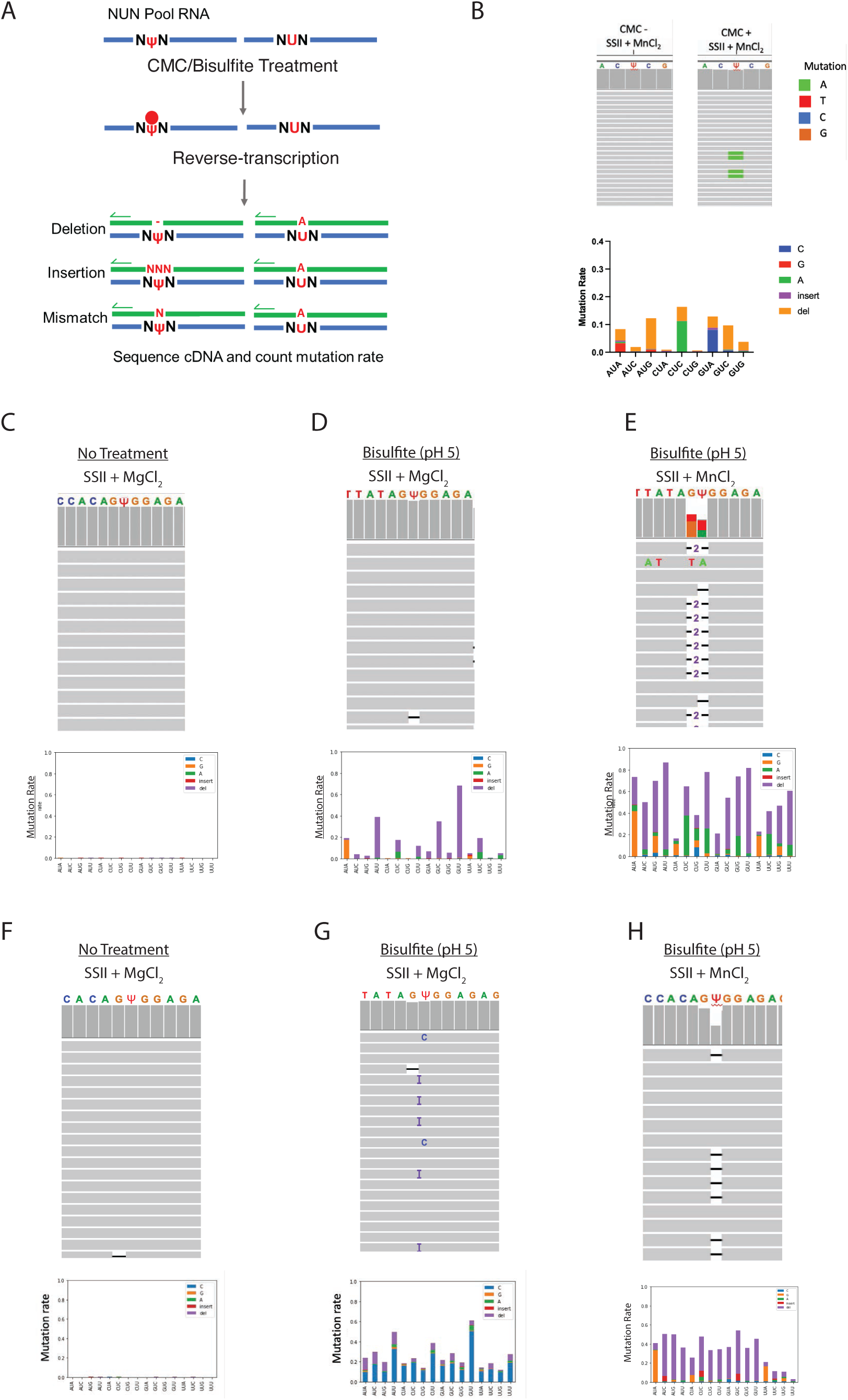
Screening of Ѱ detection methods using synthetic RNA standards. **A.** Schematic of NUN/NYN pools. RNA standards were designed with a single 100% U or 100% Ψ occupancy position, flanked by each possible adjacent nucleotide. These standards were used to screen chemical treatments and reverse-transcription conditions for sensitivity to Ψ across sequence contexts. **B-H.** Top: IGV browser view of NΨN RNA sequenced after indicated chemical treatment and reverse-transcription conditions. Bottom: Mutation rates of NΨN RNA for each sequence context at Ψ-installed position.

**Figure S2:**
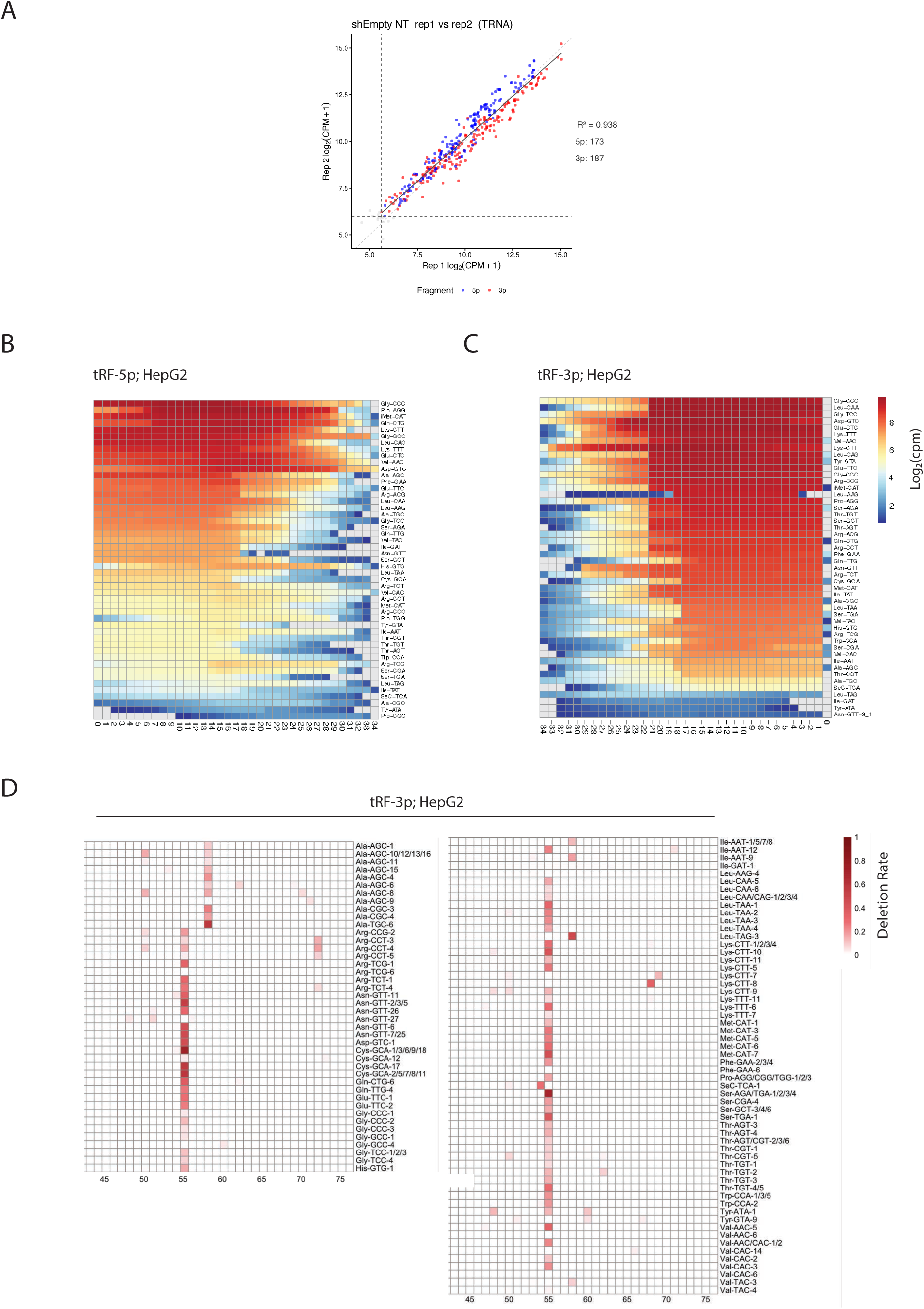
tRNA fragment bisulfite sequencing in HepG2 cells. **A.** Scatterplot of mapped tRNA fragment expression between two independent sequencing replicates. Dashed line indicates 100-read depth cutoff for Ψ site calling. **B.** IGV browser view of Glu-CTC tRF-5 after bisulfite sequencing. Deletion peak at G_7_ corresponds to a known m^2^G modification^30^. **C-D.** Coverage heatmap of most abundant 5′ and 3′ tRNA fragments from each tRNA isodecoder family. **E.** Deletion rate heatmap of 3′ tRFs after bisulfite treatment.

**Figure S3.**
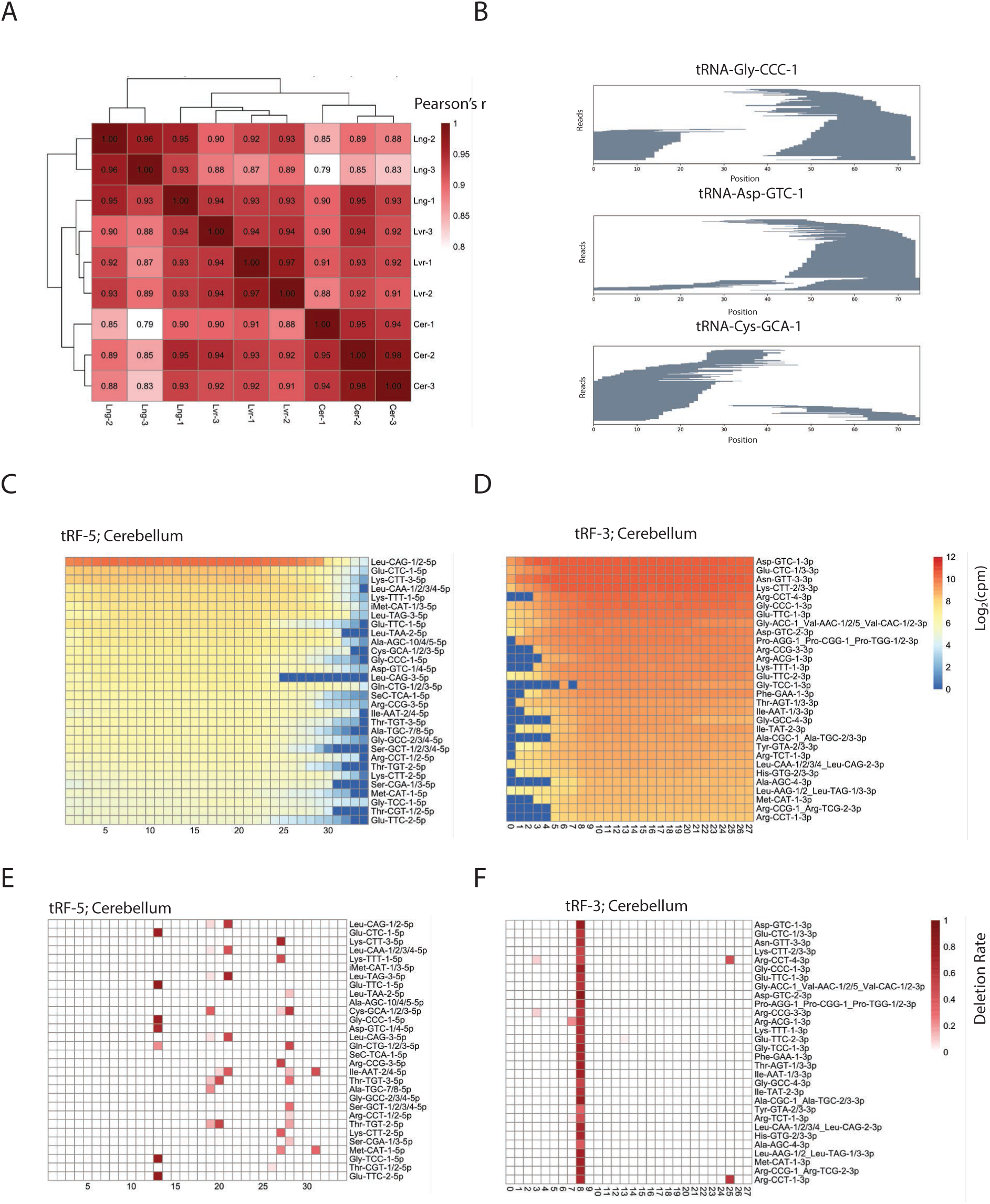
tRNA fragment bisulfite sequencing across mouse tissues. **A.** Clustered heatmap of pearson’s correlation coefficient (r) of mapped tRNA fragments across tissue replicates. **B.** Example pileups of reads mapping to full-length tRNAs in mouse cerebellum. **C-D.** Coverage heatmap of 5′ and 3′ tRNA fragments in mouse cerebellum. **E-F.** Deletion rate heatmap of 5′ and 3′ tRNA fragments in mouse cerebellum after bisulfite treatment.

**Figure S4.**
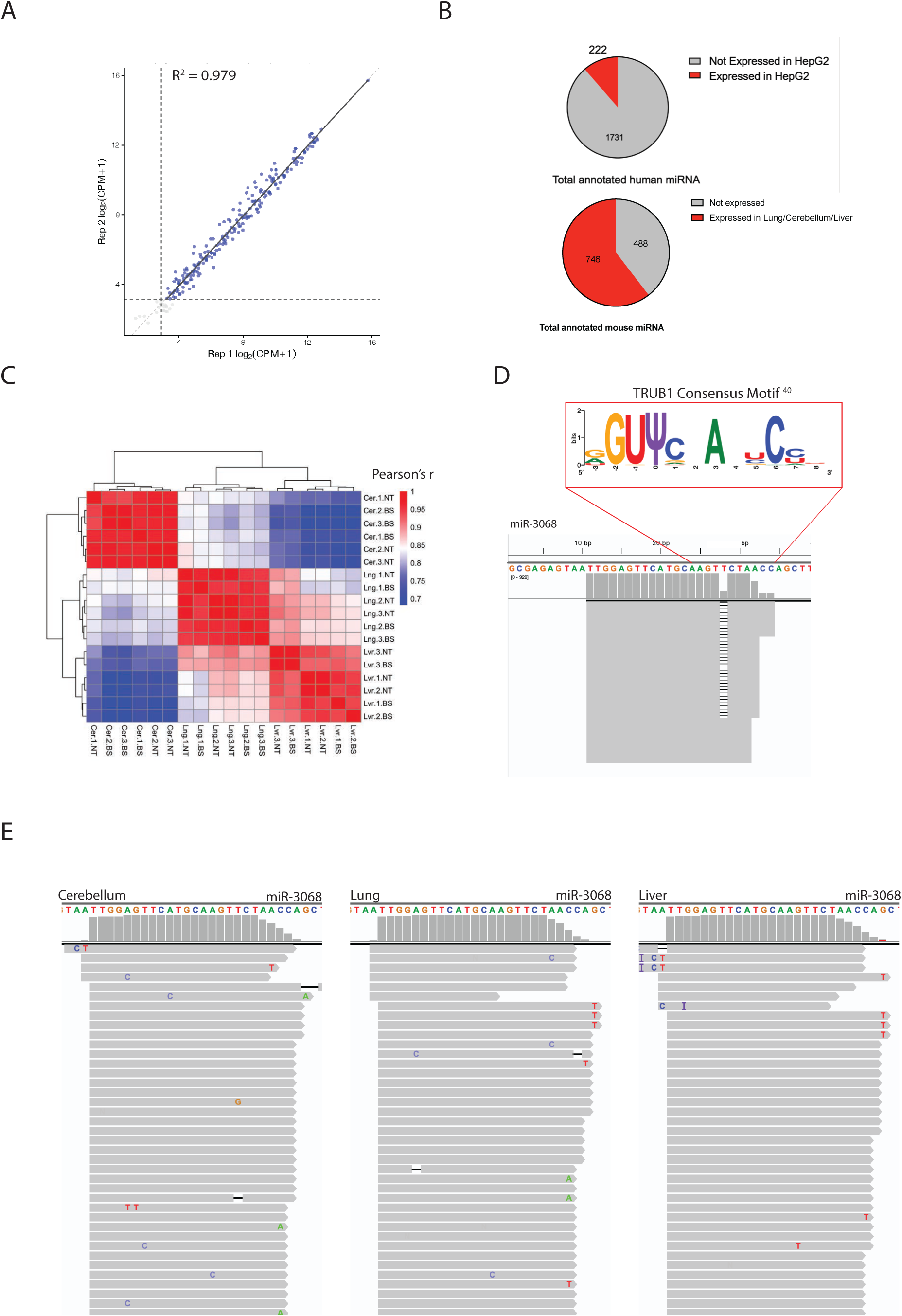
**microRNA bisulfite sequencing in HepG2 and mouse tissues**. **A.** Correlation scatterplot of mapped microRNA between two independent sequencing replicates of non-bisulfite-treated HepG2 libraries. **B.** Pie chart showing relative proportion sufficiently sequenced miRNA ( > 100 reads) in (Top) HepG2 or (Bottom) mouse tissues compared to total annotated miRNA sequences in respective species. **C.** Clustered heatmap of pearson’s correlation coefficient (r) of mapped miRNAs across tissue replicates. **D.** Graphic showing presence of Trub1 motif^40^ at miR-3068-5p Ψ position. **E.** IGV browser view of no-bisulfite miR-3068-5p sequences in mouse cerebellum, lung, and liver tissues.

**Figure S5.** RNA sequencing of miR-3068-5p transfected mouse fibroblasts. **A.** Cumulative distribution of Log_2_FC of miR3068-5p-Ψ vs Scramble-Ψ. Transcripts are binned by the strongest predicted miR-3068 site present (8 mer > 7mer-M8 > 7mer-A1 > 6mer > none) according to TargetScan. P-values calculated by Kolmogorov–Smirnov test between respective site bin and transcripts with no predicted site.

## Notes

### Competing Interest Statement

The authors have declared no competing interest.

### Funding

- National Institute of General Medical Sciences: 5R01GM101316-11

